# First record of a second invasive soft coral species, *Latissimia ningalooensis*, in southern Puerto Rico

**DOI:** 10.1101/2025.04.16.648000

**Authors:** Daniel A. Toledo-Rodriguez, Catherine S. McFadden, Nilda M. Jimenez Marrero, J. David Muñoz-Maravilla, Alex J. Veglia, Ernesto Weil, Nikolaos V. Schizas

## Abstract

Ever since the discovery of the invasive soft coral species *Xenia umbellata* (Anthozoa, Octocorallia, Malacalcyonacea, Xeniidae) on the reefs of southwestern Puerto Rico, ongoing surveys have documented its spread and potential impacts on native marine fauna. During benthic surveys conducted by scientific divers from Puerto Rico’s DNER and the Department of Marine Sciences at UPRM, colonies of xeniid soft corals were observed with morphological and color characteristics distinct from those of *X. umbellata*. Morphological and genetic barcoding analyses of four gene regions confirmed the presence of a second invasive xeniid species, *Latissimia ningalooensis*, in the southern reefs of Puerto Rico. Originally described in Western Australia, *L. ningalooensis* has recently been reported in southeastern Brazilian waters, marking its expansion into the Atlantic. The discovery of a second xeniid species in Puerto Rico following the recent introduction of *X. umbellata*, is both surprising and concerning. Highly disturbed reefs, such as those along the southern coast of Puerto Rico and the wider Caribbean, seem highly susceptible to invasive species. Recent reports of invasive soft corals and other marine species in Puerto Rico highlight the potential for some species to become a regional issue, requiring coordinated management actions across the Caribbean.

## Introduction

Invasive alien species threaten ecosystem services, *sensu* (Costanza 1997), by altering seascapes, reducing local biodiversity, and adversely affecting fishing practices, tourism, and human health (Bax et al. 2003). Caribbean coral reefs are especially vulnerable to invasive species since these coral reefs have been affected by global and local stressors during the past half-century (Hughes 1994; Eakin et al. 2010; Weil and Rogers 2011; Weil et al. 2017; Muniz-Castillo et al. 2019; Weil et al. 2019). The increasing reports of alien species in the region have raised concerns among resource managers, biologists, residents, and stakeholders across Caribbean nations. Among these, xeniid soft corals have emerged as a particularly alarming invasive taxon (Ruiz-Allais et al. 2021; Toledo-Rodriguez et al. 2024).

The first exotic xeniid soft coral reported in the Caribbean, *Unomia stolonifera* (Gohar, 1938), is native to the Indo-Pacific and was initially documented in Venezuela (Ruiz Allais et al. 2014). Since then, it has spread to other areas of the southwestern Caribbean, where it has overgrown benthic habitats, displacing sessile native fauna, particularly scleractinian corals (Ruiz Allais et al. 2014; Ruiz-Allais et al. 2021). In October 2023, another xeniid species, *Xenia umbellata* Lamarck, 1816, was detected in multiple reef locations in south and southwestern Puerto Rico, marking its first recorded presence in the Caribbean Sea (Toledo-Rodriguez et al. 2024). Both *U. stolonifera* (Espinosa et al. 2023) and *X. umbellata* (McFadden et al. unpub. data) have also been confirmed in Cuba. Native to the Red Sea, *X. umbellata* shares a concerning trait with *Unomia stolonifera*: the tendency to overgrow sessile reef organisms and substrates, potentially monopolizing entire habitats, reducing diversity, and disrupting the composition and dynamics of the invaded reef communities.

Here, we report the presence of a third invasive xeniid coral in the Caribbean Sea, *Latissimia ningalooensis* Ekins, Benayahu & McFadden, 2022, first observed in southern Puerto Rico in March 2024 (Fig. 1A). Native to northwestern Australia, *Latissimia ningalooensis* was previously reported as an invasive species in southeastern Brazil in 2017 (Mantelatto et al. 2018). The detection of three xeniid species in the Caribbean, two of which were recorded during 2023/2024, underscores the urgent need for regional collaboration to monitor new invasions and implement strategies to manage or eradicate these invasive species.

**Figure 1.**
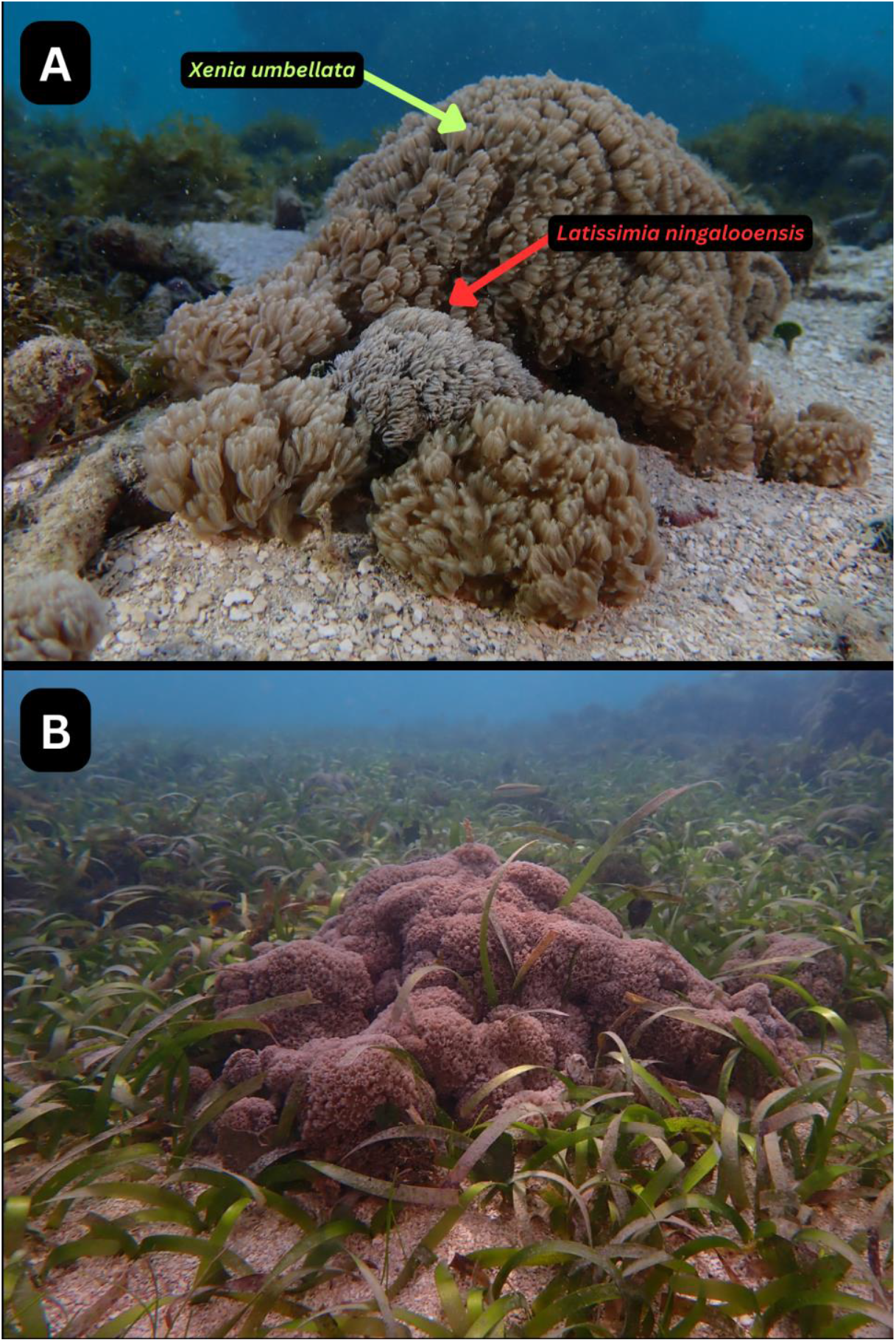
(A) Colonies of *Xenia umbellata* (green arrow) and *Latissimia ningalooensis* (red arrow) growing together on a rubble substrate near the location where the samples were collected, representing the co-existence of both xeniid species invading Caja de Muertos, southern Puerto Rico. (B) A colony of *Xenia umbellata* overgrowing a seagrass-dominated benthic habitat of *Thalassia testudinum*. Photos by Daniel A. Toledo-Rodriguez.

## Methods

### Sampling Sites

Colonies (n=2) were sampled from the northeast side of Caja de Muerto (∼17°53’47.23”N, 66°30’32.79”W), an island south of Ponce, where the presence of pulsing corals was reported. They were collected separately based on observed differences in physical characteristics. Two tissue samples were collected from each visually different colony, carefully extracted either manually or with tweezers, with the substrate, and then placed in a zipper storage bag (1 Quart). Metadata (i.e. date, depth, substrate type, and observations) were recorded. All collections were conducted as part of the implementation of the Emergency Response Strategy by personnel (NMJM) of the Puerto Rico Department of Natural and Environmental Resources to address the invasive octocorals of the Xeniidae family, which includes taxonomic validation of the invasive species. The site where the samples were collected is characterized by a fringing reef combined with patches of sand, rubble and seagrass, mainly dominated by the seagrass *Thalassia testudinum*, in which both xeniid species have been observed growing (Fig. 1). The depth range is 1 to 4 meters, with corals of the genus *Orbicella* and *Porites* being the main components of the fringing reef. This locality supports many healthy colonies of scleractinian corals. Several colonies, however, exhibit partial mortality from recent disease infections, which are now overgrown by macroalgae. Colonies of both *L. ningalooensis* and *X. umbellata* were observed overgrowing the different habitats of this reef system (Figs 1, 2).

**Figure 2.**
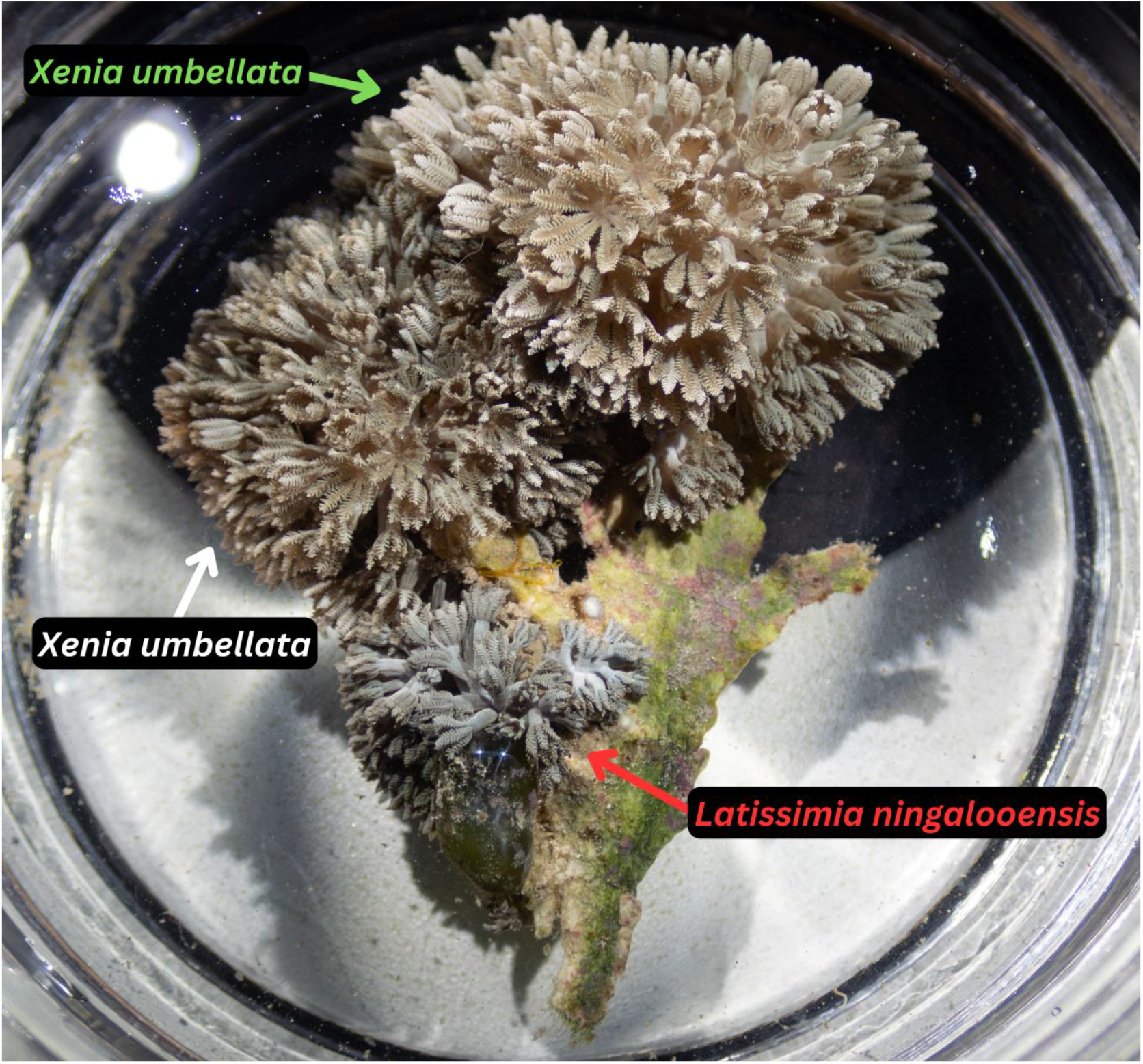
A sample of three small xeniid colonies growing on a hard substrate, (1) *Xenia umbellata* (top right), *Latissimia ningalooensis* (bottom left), and potentially another morphotype of *Xenia umbellata* (top left) showing some mild coloration differences growing on what appears to be *Udotea* spp., a seagrass. Photos by Daniel A. Toledo-Rodriguez.

### DNA procedures

We obtained DNA sequences from four genetic regions (16S/ND2, COI, mtMutS, 28S rDNA) to help us identify the new invasive xeniid. All molecular protocols used were identical to those reported in Toledo-Rodriguez et al. (2014), except that DNA traces were quality-checked and trimmed in Codon Code Aligner v. 12.0.1 (Codon Code Corp.).

### Morphological Methods

Ethanol-preserved specimens were examined under a dissecting microscope (SZX10 Olympus), where the polyp body length was measured and the number of pinnules on each side of the polyps were counted from two colonies. Sclerites were extracted from the soft coral tissue by dissolving it in 5% household bleach; sclerites were examined under light microscopy with a BX51 Olympus microscope and a (FE-SEM) Zeiss 560 VP (Fig. 3).

**Figure 3.**
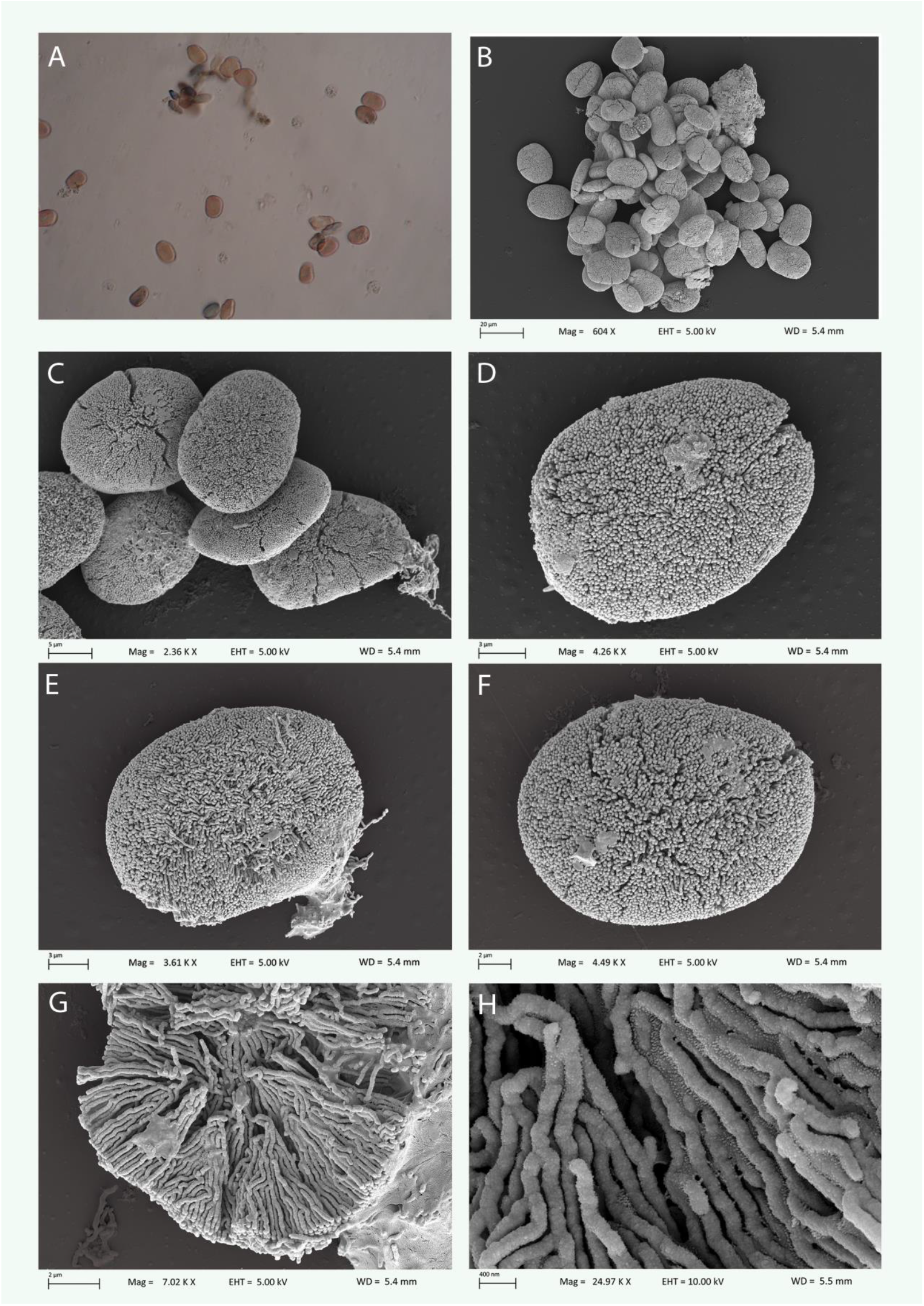
Sclerites of *Latissimia ningalooensis* A) observed under light microscopy (400X magnification with an Olympus BX51); B-H SEM photos with a (FE-SEM) Zeiss 560 VP under different magnifications, B) A clump of sclerites; C) A smaller clump of sclerites showing the ellipsoid platelet shape with a granular appearance of the sclerite surface; D, E, F) individual sclerites; G) view of the internal architecture of a damaged sclerite with a clear view of the calcite rods; H) a detailed view of the calcite rods that make up the sclerites of *L. ningalooensis*.

## Results and Discussion

Morphology: Colonies of *Latissimia ningalooensis* were observed in Caja de Muertos, Puerto Rico, displaying a range of color variations, including bluish hues, dark browns, and grayish tones (Fig. 2). Colonies exhibited encrusting morphology with polyps arising from a spreading thick membrane (∼1 mm). The non-retractile polyps had tentacles that ranged from 1.9 – 2.8 mm in length, with 2 – 3 rows of short, stubby pinnules on each side. Measurements from the genetically confirmed colony and the second *L. ningalooensis* colony, showed pinnules on either side of their tentacles to range from 17 – 37 and 15 – 37, respectively. Pinnule sizes ranged from 0.3 mm – 0.4 mm in length and 0.1 mm – 0.2 mm in width, with closely juxtaposed arrangements.

Sclerites have a typical ellipsoid platelet-like morphology (Fig. 3) with slight variation in shape (Fig. 3B-F), found in high numbers throughout the polyps. Very few sclerites (Fig 3B) are observed with narrow waists, as also documented by (Benayahu et al. 2022). Size varies from 14 μm x 18 μm to 15 μm x 19 μm, based on measurements on sclerites in Figs 3D, E, and F. Benayahu et al. (2022) report a slightly wider size range of sclerites from the Australian soft corals (11 μm x 13 μm to 16 μm x 18 μm) and (Mantellato et al. 2018) a slightly longer sclerite diameter (∼ 20 μm). The surface of sclerites has a spongy, granular appearance, only visible under SEM observations. Sclerites are composed of long, continuous calcite rods oriented perpendicular to the sclerite surface (Fig. 3G, H). Overall, the shape, dimensions, surface texture, and micromorphology of sclerites from the Puerto Rican colonies are very similar to the sclerite description of the type specimens of *L. ningalooensis* from Australia (see Fig. 4 (Benayahu et al. 2022).

The colony morphology of *L. ningalooensis* colonies in Puerto Rico is similar to those reported in Australia and Brazil: encrusting, featuring a spreading membrane and non-retractile polyps. There are some gross colony morphological differences between the type material from Australia and the material from Brazil and Puerto Rico, including the polyp body length and the number of pinnules on the tentacles. The polyp body length was 3.5 mm in Australian colonies (Benayahu et al. 2022) but varied by an order of magnitude in Brazilian colonies, from 3.5 mm to 3.75 cm (Mantellato et al. 2018), likely due to differing degrees of contraction. In Puerto Rico, the polyp body length of ethanol-preserved specimens ranges from 1.2 mm – 4.9 mm in length and 1 mm – 3.6 mm in width. The polyps of Puerto Rican *L. ningalooensis* have rather similar numbers of pinnules to those observed in the holotype specimen from western Australia (two rows of only 18–22 pinnules) but fewer than the Brazilian colonies, which have two rows of 35–50 pinnules on either side of the tentacle. The high variation of pinnule counts within and between soft coral colonies renders this character unreliable for xeniid species delineation (Benayahu et al. 2022).

DNA barcoding: The length of DNA sequences after end trimming was reduced to 690bp (16S/ND2), 844bp (COI), 715bp (mtMutS), and 773bp (28S). BLASTn searches of the four genes revealed 100% sequence identity with the species *Latissimia ningalooensis* (e.g., GenBank Accession Numbers, 16S/ND2: XXXXX; COI: XXXXX; mtMutS: PV172462; 28S: PQ874183). In addition, sequences of all four genes were compared against other published (Benayahu et al. 2022) and unpublished xeniid sequences (McFadden et al, mss. in review), verifying the morphological species diagnosis.

The Caribbean Sea has long been characterized as a disease hotspot (e.g. (Weil 2004; Weil and Rogers 2011; van Woesik and Randall 2017; Weil et al. 2019). However, recent reports of the presence and spread of at least three invasive and aggressive xeniid octocorals—*Unomia stolonifera* (e.g., Venezuela), *Xenia umbellata* (e.g., Puerto Rico and Cuba), and *Latissimia ningalooensis* (Puerto Rico)— suggest that the region may also be emerging as a hotspot for invasive soft coral species. The type locality of *Latissimia ningalooensis* is Ningaloo Reef, Western Australia, but it has also been recorded in other reef localities across northwestern Australia and the Northern Territory (Darwin). In its native range, live colonies of *L. ningalooensis* exhibit a light brown coloration with a variable blue tinge that resemble colors observed in Puerto Rico in addition to dark brown with grayish tones (Figs 1, 2).

This species was first reported in the Atlantic Ocean in southeastern Brazil, where it was initially identified as *Sansibia* sp. before *L. ningalooensis* was formally recognized as a distinct genus and species (Benayahu et al. 2022). *Latissimia ningalooensis* colonies were detected on September 18, 2017, on shallow subtidal tropical rocky reefs at Ilha Grande Bay, Rio de Janeiro State, southeast Brazil (Mantelatto et al. 2018). They have been observed at all depths on hard substrata, including rocks, metal, plastic, and even tires, extending to the shore-sand interface (Mantelatto et al. 2018; de Carvalho-Junior et al. 2023), and significantly changing the species composition between invaded and control areas (Pires-Teixeira et al. 2024). An increase in the abundance of *L. ningalooensis* in shallow areas indicates the species’ potential to spread outside the inner part of Ilha Grande Bay, where the colonies were first detected (Pires-Teixeira et al. 2024). Their presence is regarded as threatening to the structure and functioning of macroalgal-dominated rocky reefs (de Carvalho-Junior et al. 2023). The co-occurrence of *L. ningalooensis* and *Xenia umbellata* in the southern reef region of Puerto Rico suggests a common origin of introduction for these invasive species. The most common hypothesis is that since both octocorals are common in the marine aquarium trade, they could have been released accidentally or intentionally by marine aquarists as appears to be the case in Brazil (Mantellato et al. 2018). However, other pathways of introductions, such as commercial shipping or transport via floating natural and artificial substrata (Hoeksema et al. 2023) cannot be ruled out. Efforts to eradicate *L. ningalooensis* in Brazil have so far proven unsuccessful, indicating that this species is a persistent invader capable of altering reef ecosystems, similar to *Unomia stolonifera* in Venezuela (Ruiz Allais et al. 2014; Ruiz-Allais et al. 2021), *Xenia umbellata* in Puerto Rico (e.g. Fig. 2B and Toledo-Rodriguez et al. pers. obs) and *Sarcothelia* sp. in Brazil (Lolis et al. 2023).

Frequent reef surveys and rapid identification of alien species remain the best line of defense against such invasions. Catching the “first colony” is probably the only way of eradicating the invader. Once the colony is established and starts to reproduce asexually (clones, fragmentation, etc.) or sexually, it becomes almost impossible to eliminate. An important and time-sensitive study involves characterizing the reproductive biology and dispersal capabilities of these three species, as this will inform optimal containment or eradication practices.

The ongoing and significant loss of live cover among scleractinian corals, octocorals, colonial zoanthids, sponges, crustose coralline algae, and other organisms across Caribbean coral reef ecosystems is driven by the emergence of new infectious diseases, widespread bleaching events, and human-induced deterioration of local and regional environmental conditions. This has resulted in vast areas of bare substrate (Weil and Rogers 2011; Weil et al. 2017; McClanahan et al. 2018; Weil et al. 2019), creating ideal conditions for the establishment, survival and spread of these highly damaging exotic octocoral species.

Collaborative initiatives of continuous observations and surveys of potential spread and dispersion routes, as well as interactions with local benthic fauna, should be supported by well-funded programs to achieve effective management/eradication of invasive species. As an example, DNER of Puerto Rico is collaborating with academia to further the genetic analysis of invasive soft corals as a tool to decipher propagation pathways. Coordinated efforts involving biologists, concerned citizens (e.g., divers), stakeholders (e.g., fishers), and resource management agencies are essential to effectively address the growing threat of invasive species in the Caribbean.

## Acknowledgments

We thank the Department of Marine Sciences (UPRM), the Department of Natural and Environmental Resources of Puerto Rico and Blue Water Scuba for providing field and logistical support. We appreciate Geli Boutou’s help in editing and arranging Fig 3. We acknowledge the Department of Natural and Environmental Resources for granting the sampling permit (O-VS-PVS15-SJ-01444-14032024) used in this study.

## Author contributions

Daniel A. Toledo-Rodriguez, Nilda M. Jimenez, Nikolaos V. Schizas, and Catherine S. McFadden conceived and developed the study. Nilda M. Jimenez and Daniel A. Toledo-Rodriguez provided the samples. Daniel A. Toledo-Rodriguez produced the genetic data and Nikolaos V. Schizas and Catherine S. McFadden analyzed the genetic data. Nikolaos V. Schizas and J. David Muñoz-Maravilla produced and analyzed the morphological data. All authors contributed to writing the manuscript.

## Data availability

The DNA sequences referred to in this manuscript have been deposited in GenBank (Accession Numbers, 16S/ND2: XXXX; COI: XXXX; mtMutS: PV172462; 28S: PQ874183).

## Declarations

## References

Bax N, Williamson A, Aguero M, Gonzalez E, Geeves W (2003) Marine invasive alien species: a threat to global biodiversity. Marine Policy 27: 313–323 doi 10.1016/s0308-597x(03)00041-1

Benayahu Y, Ekins M, Ofwegen LPV, Samimi-Namin K, McFadden CS (2022) On some encrusting Xeniidae (Octocorallia): Re-examination of the type material of Sansibia flava (May, 1898) and a description of new taxa. Zootaxa 5093: 421–444 doi 10.11646/zootaxa.5093.4.3.

Costanza R, d’Arge, R., de Groot, R. et al. (1997) The value of the world’s ecosystem services and natural capital. Nature 387: 253–260 doi 10.1038/387253a0

de Carvalho-Junior L, Neves LM, Teixeira-Neves TP, Cardoso SJ, Alicea-Rodríguez L (2023) Long-term changes in benthic communities following the invasion by an alien octocoral in the Southwest Atlantic, Brazil. Marine Pollution Bulletin 186: 114386 doi 10.1016/j.marpolbul.2022.114386

Eakin CM, Morgan JA, Heron SF, Smith TB, Liu G, Alvarez-Filip L, Baca B, Bartels E, Bastidas C, Bouchon C, Brandt M, Bruckner AW, Bunkley-Williams L, Cameron A, Causey BD, Chiappone M, Christensen TR, Crabbe MJ, Day O, de la Guardia E, Diaz-Pulido G, DiResta D, Gil-Agudelo DL, Gilliam DS, Ginsburg RN, Gore S, Guzman HM, Hendee JC, Hernandez-Delgado EA, Husain E, Jeffrey CF, Jones RJ, Jordan-Dahlgren E, Kaufman LS, Kline DI, Kramer PA, Lang JC, Lirman D, Mallela J, Manfrino C, Marechal JP, Marks K, Mihaly J, Miller WJ, Mueller EM, Muller EM, Orozco Toro CA, Oxenford HA, Ponce-Taylor D, Quinn N, Ritchie KB, Rodriguez S, Ramirez AR, Romano S, Samhouri JF, Sanchez JA, Schmahl GP, Shank BV, Skirving WJ, Steiner SC, Villamizar E, Walsh SM, Walter C, Weil E, Williams EH, Roberson KW, Yusuf Y (2010) Caribbean corals in crisis: record thermal stress, bleaching, and mortality in 2005. PLoS One 5: e13969 doi 10.1371/journal.pone.0013969

Espinosa J, Estrada R, Ruiz-Allais JP (2023) Presencia en Cuba de la especie marina invasora Unomia stolonifera (Gohar, 1938) (Octocorallia, Alcyonacea). Acciones para su control y eliminación. Revista Investigaciones Marinas 43: 140–146 doi 10.5281/zenodo.8018890

Hoeksema BW, Samimi-Namin K, McFadden CS, Rocha RM, van Ofwegen LP, Hiemstra AF, Vermeij MJA (2023) Non-native coral species dominate the fouling community on a semi-submersible platform in the southern Caribbean. Mar Pollut Bull 194: 115354 doi 10.1016/j.marpolbul.2023.115354

Hughes TP (1994) Catastrophes, Phase Shifts, and Large-Scale Degradation of a Caribbean Coral Reef. Science 265: 1547–1551 doi 10.1126/science.265.5178.1547

Lolis LA, Miranda RJ, Barros F (2023) The effects of an invasive soft coral on the structure of native benthic communities. Marine Environmental Research 183: 105802 doi 10.1016/j.marenvres.2022.105802

Mantelatto MC, Silva AGD, Louzada TDS, McFadden CS, Creed JC (2018) Invasion of aquarium origin soft corals on a tropical rocky reef in the southwest Atlantic, Brazil. Marine Pollution Bulletin 130: 84–94 doi 10.1016/j.marpolbul.2018.03.014

McClanahan TR, Weil E, Baird AH (2018) Consequences of Coral Bleaching for Sessile Reef Organisms. In: van Oppen M, Lough J (eds) Coral Bleaching. Springer, Cham, pp 231–263

Muniz-Castillo AI, Rivera-Sosa A, Chollett I, Eakin CM, Andrade-Gomez L, McField M, Arias-Gonzalez JE (2019) Three decades of heat stress exposure in Caribbean coral reefs: a new regional delineation to enhance conservation. Scientific reports 9: 11013 doi 10.1038/s41598-019-47307-0

Pires-Teixeira LM, Neres-Lima V, Creed JC (2024) Diversity analysis and trophic structure of a recently invaded tropical rocky shore. Biological Invasions 27 doi 10.1007/s10530-024-03478-0

Ruiz Allais JP, Amaro ME, McFadden CS, Halász A, Benayahu Y (2014) The first incidence of an alien soft coral of the family Xeniidae in the Caribbean, an invasion in eastern Venezuelan coral communities. Coral Reefs 33: 287–287 doi 10.1007/s00338-013-1122-1

Ruiz-Allais JP, Benayahu Y, Lasso-Alcalá OM (2021) The invasive octocoral Unomia stolonifera (Alcyonacea, Xeniidae) is dominating the benthos in the Southeastern Caribbean Sea. Memoria de la Fundación La Salle de Ciencias Naturales 79: 63–80 doi 10.5281/ZENODO.4784709

Toledo-Rodriguez DA, Veglia A, Jimenez Marrero NM, Gomez-Samot JM, McFadden CS, Weil E, Schizas NV (2024) Shadows over Caribbean reefs: Identification of a new invasive soft coral species, Xenia umbellata, in southwest Puerto Rico. bioRxiv doi 10.1101/2024.05.07.592775

van Woesik R, Randall CJ (2017) Coral disease hotspots in the Caribbean. Ecosphere 8(5):e01814.10.1002/ecs2.1814 doi 10.1002/ecs2.1814

Weil E (2004) Coral Reef Diseases in the Wider Caribbean. In: Rosenberg E, Loya Y (eds) Coral Health and Disease. Springer, Berlin, Heidelberg pp 35–68

Weil E, Hernández-Delgado EA, Gonzalez M, Williams S, Suleimán-Ramos S, Figuerola M, Metz-Estrella T (2019) Spread of the New Coral Disease “SCTLD” into the Caribbean: Implications for Puerto Rico Reef Encounters 34: 38–43

Weil E, Rogers CS (2011) Coral Reef Diseases in the Atlantic-Caribbean. In: Dubinsky Z, Stambler N (eds) Coral Reefs: An Ecosystem in Transition. Springer, Dordrecht, pp 465–491

Weil E, Rogers CS, Croquer A (2017) Octocoral Diseases in a Changing Ocean. In: Rossi S, Bramanti L, Gori A, Orejas C (eds) Marine animal forests: The ecology of benthic biodiversity hotspots. Springer, pp 1–55

